# Physiological and pathological brain activation in the anesthetized rat produces hemodynamic-dependent cortical temperature increases that can confound the BOLD fMRI signal

**DOI:** 10.1101/205609

**Authors:** Samuel S. Harris, Luke W. Boorman, Devashish Das, Aneurin J. Kennerley, Paul S. Sharp, Chris Martin, Peter Redgrave, Theodore H. Schwartz, Jason Berwick

## Abstract

Anesthetized rodent models are ubiquitous in pre-clinical neuroimaging studies. However, because the associated cerebral morphology and experimental methodology results in a profound negative brain-core temperature differential, cerebral temperature changes during functional activation are likely to be principally driven by local inflow of fresh, core-temperature, blood. This presents a confound to the interpretation of blood-oxygenation level-dependent (BOLD) functional magnetic resonance imaging (fMRI) data acquired from such models, since this signal is also critically temperature-dependent. Nevertheless, previous investigation on the subject is surprisingly sparse. Here, we address this issue through use of a novel multi-modal methodology in the urethane anesthetized rat. We reveal that sensory stimulation, hypercapnia and recurrent acute seizures induce significant increases in cortical temperature that are preferentially correlated to changes in total hemoglobin concentration, relative to cerebral blood flow and oxidative metabolism. Furthermore, using a phantom-based evaluation of the effect of such temperature changes on the BOLD fMRI signal, we demonstrate a robust inverse relationship between the two. These findings indicate that temperature increases, due to functional hyperemia, should be accounted for to ensure accurate interpretation of BOLD fMRI signals in pre-clinical neuroimaging studies.

## Introduction

Negative brain-core temperature differentials, such that the brain temperature is lower than that of the body-core, are commonly seen in small animals whose brains have high surface to volume ratios and are particularly vulnerable to heat exchange with the environment (Zhu et al., 2009). Anesthesia can amplify these effects (Shirey et al., 2015), insofar as brain temperatures in anesthetized rats are approximately 1°C lower in the deep brain than in the core, and decrease progressively outward to a maximal deficit of 3-4°C at the cortical surface (Zhu et al., 2006). Cortical temperatures directly below craniotomies in isoflurane-anesthetized mice can be even more dramatic, being up to 10°C cooler than the core body temperature (Kalmbach and Waters, 2012), possibly as a result of a profound ‘toxic insult’ response to this anesthetic agent (Shirey et al., 2015). Inflow of fresh arterial blood at core temperature into functionally-activated cortical regions, in both anesthetized and awake rodents, thus leads to a large increase in local tissue temperature through heat exchange (Zhu et al., 2009). Heat production during increases in cerebral metabolic rate of oxygen (CMRO_2_) with functional activation also contribute to changes in local brain temperature, but are thought to have a comparatively small influence on brain temperature relative to blood flow in the presence of large resting brain/core-body temperature differentials (Yablonskiy et al., 2000; Collins et al., 2004; Zhu et al., 2006; Zhu et al., 2009). Nevertheless, this has not been comprehensively studied across a range of physiological and pathological activation.

Changes in brain temperature also affect the rate of local chemical reactions, such that the average Q10 temperature coefficient (change in chemical reaction rate with a temperature increase of 10°C) in the brain is 2.3 (Swan, 1974), suggesting that a small change in brain temperature could induce a significant change in cerebral metabolic rate (Zhu et al., 2009). Furthermore, brain temperature changes can also affect the affinity of hemoglobin for oxygen (the Bohr effect), with blood oxygen saturation changing by several percent with every °C change (Yablonskiy et al., 2000). These phenomena have been exploited to infer changes in brain temperature from human blood-oxygenation level dependent (BOLD) functional magnetic resonance (fMRI) signals (Yablonskiy et al., 2000; Sotero and Iturria-Medina, 2011), which rely on the incompletely understood interplay between cerebral blood flow, metabolism and hemoglobin oxygenation (Ogawa et al., 1993). Decreased oxygen affinity of hemoglobin has also been suggested to underpin a negative correlation between BOLD signal intensity and manipulated increases in body temperature (37-39°C) of anesthetized rats (Vanhoutte et al., 2006). Importantly, however, MR relaxation properties were also found to change with induced temperature changes in human adipose tissue, muscle, and bone samples (Petrén-Mallmin et al., 1993). This implies that BOLD fMRI signals during functional activation can be modulated by non-physiological temperature variations in the absence of hemoglobin or metabolic fluctuations. Remarkably, this confound has not been previously accounted for in BOLD neuroimaging studies using anesthetized rodent models, which possess profound negative brain-core temperature differentials and are thus predisposed to large increases in brain temperature during functional hyperemia (Zhu et al., 2009). Importantly, this effect may also have implications for pre-clinical studies using craniotomies to elucidate the underpinnings of the BOLD signal through novel combination with other optical modalities such as optical imaging spectroscopy (Kennerley et al., 2005), calcium imaging (Schulz et al., 2012; Liang et al., 2017), optogenetics (Lee et al., 2010; Schmid et al., 2017) and, potentially, two-photon microscopy (Cui et al., 2017).

Accordingly, we have sought to address these unresolved questions through development of a novel multi-modal methodology that provides concurrent recordings of cortical temperature, blood flow and total hemoglobin concentration, as well as estimates of CMRO_2_ through measures of tissue oxygenation. We aimed to quantify and interrogate the relationship between cortical temperature responses and hemodynamic and metabolic variables during sensory stimulation, hypercapnia, and recurrent acute seizures, in the urethane-anesthetized rat cortex - a popular model employed in pre-clinical BOLD fMRI studies. We demonstrate that this spectrum of brain activation induces significant increases in cortical temperature that are most closely correlated to changes in total hemoglobin concentration. Finally, through phantom-based evaluation, we show that such temperature increases, in of them themselves, markedly, and inversely, affect the BOLD fMRI signal. Our results provide important insights into the mechanisms underlying temperature increases during functional activation in a common pre-clinical model, and underscore the importance of considering temperature-dependent confounds in relevant neuroimaging studies.

## Methods

### Animal preparation and surgery

All procedures were conducted with approval from the UK Home Office under the Animals (Scientific Procedures) Act of 1986. Female hooded Lister rats (total N=11 weighing 260-325g) were kept in a 12-hr dark/light cycle environment at a temperature of 22°C, with food and water *ad libitum*. Animals were anesthetized with urethane (1.25 g/kg, i.p). Urethane is a popular anesthetic in non-recovery pre-clinical neuroimaging studies as it provides long-term stable anesthesia reminiscent of natural sleep (Pagliardini et al., 2013), and preserves excitatory and inhibitory synaptic transmission (Sceniak and MacIver, 2006) and neurovascular coupling (Berwick et al., 2008). Furthermore, the spatial–temporal pattern of stimulus induced hemodynamic responses (Devor et al., 2005), and the relationship between neural activity and the BOLD fMRI response (Huttunen et al., 2008), do not differ between urethane and alpha-chloralose, another anesthetic widely used in neurovascular studies (Masamoto and Kanno, 2012). Animal core-body temperature was maintained at a stable 37°C using a homoeothermic blanket and rectal probe (Harvard Apparatus) and room temperature was thermostatically controlled to be a constant at 19.1°C. Animals were tracheotomized and artificially ventilated with medical air, with blood-gas and end-tidal CO_2_ measurements taken to adjust ventilator parameters and maintain the animal within normal physiological limits. The left femoral artery and vein were cannulated to allow the measurement of arterial blood pressure and phenylephrine infusion (0.13-0.26 mg/h to maintain normotension between 100 and 110 mmHg), respectively (Berwick et al., 2005; Berwick et al., 2008). The animal was secured in a stereotaxic frame throughout experimentation and the skull overlying the right parietal cortex thinned to translucency for optical imaging. A layer of cyanoacrylate glue was applied over the thinned skull preserve skull integrity and reduce optical specularities from the brain surface.

### Optical Imaging spectroscopy and localization of whisker barrel cortex

Two-dimensional optical imaging spectroscopy (2D-OIS) was used to produce images over time of total hemoglobin (Hbt) concentration, using a heterogeneous tissue model as described previously (Berwick et al., 2005; Berwick et al., 2008; Kennerley et al., 2009). Hbt can be further interpreted as CBV, under the reasonable assumption of a constant hematocrit. Briefly, the cortex was illuminated at four wavelengths (495±31 nm, 559±16 nm, 575±14 nm and 587±9 nm FWHM) using a Lambda DG-4 high speed filter changer (Sutter Instrument Company, Novata, CA, USA) and image data synchronized to the filter switching and recorded at 8Hz using a Dalsa 1M30P camera (Billerica, MA, USA, each pixel representing ~75μm^2^). Spectral analysis of image data consisted of a path length scaling algorithm (PLSA), which consisted of a modified Beer–Lambert law in conjunction with a path-length correction factor for each wavelength used, based on multi-layered Monte Carlo simulations of light transport through tissue (Wang et al., 1995; Berwick et al., 2005; Berwick et al., 2008). The 2D-OIS camera lens was fitted with a 750nm low-pass filter to prevent cross-talk from a Laser-Doppler flowmetry (LDF) probe operating at 830±10nm (see below), and the transmission curve of the filter accounted for in the spectral analysis. The somatosensory barrel cortex was localized prior to experiments by briefly electrically stimulating the left (contralateral) whisker pad using two subcutaneous electrodes (30 trials, 2s, 5Hz, 1.2mA, 0.3ms pulse width). Resultant 2D-OIS data were averaged and subjected to the aforementioned spectral analysis, and spatiotemporal changes in Hbt were analyzed using statistical parametric mapping (SPM) so as to localize the cortical region activated by whisker stimulation (as described previously, Harris et al., 2013) (Figure 1A).

### Multi-sensor probe recordings

A multi-modal sensor comprising of a thermocouple, luminescence-based oxygen, and LDF probe (Oxford Optronix, UK) was used to record cortical temperature, tissue oxygenation (tpO_2_) and blood flow (CBF), respectively, during functional activation (Trübel et al., 2006). The sensor was implanted into barrel cortex after localization of the region by 2D-OIS. The thermocouple and luminescence probes provided an absolute measure of temperature (±0.1°C) and tpO_2_ (±0.1mmHg), respectively, with the laser Doppler probe providing a measure of relative changes in CBF. Signals were amplified and digitized using a 2-channel OxyFlo Pro system (Oxford Optronix, UK) and sampled at 1Hz using a CED Power 1401 and Spike2 software (Cambridge Electronic Design, Cambridge, UK). The multi-sensor probe was affixed to a stereotaxic holder and inserted to a cortical depth of 500μm.

### Brain activation paradigm

Physiological and pathological functional activation was induced by somatosensory stimulation, graded hypercapnia and induction of recurrent acute focal neocortical seizures. Whisker stimulation consisted of 16-second trains of electrical pulses (5 Hz, 1.2 mA intensity, 0.3ms pulse width) delivered to the whisker pad using two subcutaneous electrodes (15 trials, 70s inter-trial interval, 10-second baseline, 5 animals) (Harris et al., 2013). Hypercapnia was induced in two further independent experimental runs, during which the fraction of inspired CO_2_ (FiCO_2_) was increased to 5% or 10% for 190s (3 trials, 380s inter-trial interval, 20s baseline, 5 animals). Experimental seizures were induced in a separate group of animals (N=6) since hypercapnia has been previously shown to have a strong anti-convulsant action (Tolner et al., 2011). Fluidic probes were loaded with the potassium channel blocker 4-aminopyridine (4-AP, 15mM, 1μl, Sigma, UK) to induce recurrent focal neocortical ictal-like discharges, as described in detail previously (Harris et al., 2014a; Ma et al., 2014). Briefly, a 10μl Hamilton syringe and syringe pump (World Precision Instruments Inc., FL, USA) was used to infuse 4-AP into right barrel cortex at a depth of 1500μm over a 5-minute period (0.2μl/min) (Harris et al., 2014a; Harris et al., 2014b). Multi-sensor and 2D-OIS measures were recorded concurrently for 7000s with 4-AP injection following a 280s baseline period.

### fMRI phantom experiments

Phantom BOLD-fMRI was conducted using a 7 Tesla preclinical MRI system (Bruker BioSpec, 310mm bore). The phantom consisted of a sterile polypropylene tube (length 114mm × diameter 28mm, Sarstedt AG & Co, Germany) filled with saline (50ml, 0.9%). An MR-compatible thermistor probe (YSI-402, CWE, USA), connected to a temperature monitor (±0.1°C, TC-100, CWE, USA), was placed at the center of the phantom, and phantom temperature continuously sampled throughout fMRI experiments at 100Hz using a CED Power 1401 and Spike2 software (Cambridge Electronic Design, Cambridge, UK). The phantom was preheated prior to experiments to a temperature of 40°C using a homoeothermic blanket (Harvard Apparatus), placed inside the magnet bore, and allowed to cool. BOLD-fMRI data were acquired during phantom cooling using a fast low-angle shot (FLASH) sequence with flip angle 30°, field of view 35×35mm, acquisition matrix 128 × 128, slice thickness 2mm, and TR/TE = 125ms/6ms, 125ms/12ms or 1000ms/6ms (number of acquisitions = 250, 250 and 30), respectively. A 35mm ID volume resonator was used for RF transmission and reception. A region of interest (ROI) on resultant coronal images, imported into Matlab (MathWorks, USA), was selected adjacent to the thermistor location, and the signal intensity over time averaged across selected voxels. Changes in phantom temperature within the range 38-24°C during each BOLD fMRI acquisition were subsequently extracted and related to concomitant BOLD signal intensity.

### Data and statistical analysis

Cerebral metabolic rate of oxygen (CMRO_2_) was estimated from CBF and tissue oxygenation (tpO_2_) recordings using the following relationship (Gjedde, 2005):

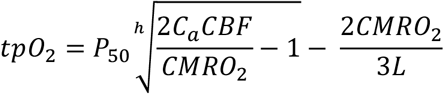

where tpO_2_ is the measured tissue oxygen tension. *P*_*50*_ and *h* denote the half-saturation tension (36mmHg) and Hill coefficient of the oxygen–hemoglobin dissociation curve (2.7), and *C*_*a*_ is the arterial oxygen concentration (8μM/ml). *L* is the effective diffusion coefficient of oxygen in brain tissue and was calculated to be 7.47±0.91μM/100g/min/mmHg (N=21, across stimulation, hypercapnia and seizure experiments) based on trial-averaged baseline tpO_2_ values in each experiment, and literature values of baseline CBF (47ml/100g/min) (He et al., 2007) and CMRO_2_ (219μM/100g/min) (Zhu et al., 2002).

Hbt time-series responses in barrel cortex to whisker stimulation were extracted by defining a ROI over Hbt image data in each animal, such that pixels possessing >80% of the maximum change in evoked Hbt during the stimulation period were automatically selected. This same ROI was also employed for extraction of Hbt time-series responses to graded hypercapnia in the same animal. Due to variable propagation dynamics during seizure activity (Ma et al., 2012), we manually defined a ROI which flanked the multi-sensor and fluidic probes, and encapsulated a well-mixed vascular compartment (arterial, venous and parenchymal). Hbt time-series of all pixels in each ROI were subsequently averaged for each animal and normalized to the baseline period prior to sensory stimulation, hypercapnia or 4-AP infusion.

Trial-by-trial analysis of the relationship between temperature and hemodynamic-metabolic variables during stimulation, hypercapnia and seizures, were performed after standardization of the data using z-scores. Timeseries of all variables are presented as a percentage change from baseline, and errors are given as the standard error from the mean (SEM). Regression models were fitted using non-linear ordinary least squares. Wilcoxon signed rank tests were used to test significant changes in individual variables, and significant differences between conditions assessed using a Kruskal-Wallis test with Tukey-Kramer correction for multiple comparisons.

The focus of the present paper is the examination of cortical temperature changes during a range of functional activation types, and how these relate to hemodynamic-metabolic changes. Specific emphasis is given to the effects of these temperature changes, *per se*, on the BOLD fMRI signal. Tissue oxygenation and laminar neural activity dynamics in the context of neural-hemodynamic coupling, have been investigated recently, educed using the same multi-modal methodology described herein, in our companion paper (Harris et al., 2017).

## Results

### Temperature increases during a range of brain activations are Hbt-dependent

We exploited our methodology to investigate cortical temperature changes, and their hemodynamic-metabolic underpinnings, during stimulation, hypercapnia and recurrent seizures. Baseline cortical temperature across animals prior to experiments was 32.4±0.3°C (N=11) while core/body temperature was maintained at 37 °C. This temperature is consistent with upper/middle neocortical laminae in anesthetized rats under controlled conditions being characteristically 3-4°C cooler than core body temperature (Zhu et al., 2006). Cortical temperature increased significantly during sensory stimulation (peak change in temperature: 0.4±0.03°C, p<0.05, 1-tailed Wilcoxon signed rank test), hypercapnia (peak change in temperature: 0.9±0.1°C and 1.6±0.2°C, 5% and 10% FICO_2_ respectively, N=5 in both cases, p<0.05, 1-tailed Wilcoxon signed rank test) and recurrent seizures (peak change in temperature: 1.8±0.1°C, N=6, p<0.05, 1-tailed Wilcoxon signed rank test) (Figure 1B). Notably, significant differences in peak temperature were only observed between whisker stimulation and 10% hypercapnia, and whisker stimulation and recurrent seizures (p<0.05 and p<0.01, respectively, Kruskal-Wallis test with Tukey-Kramer correction for multiple comparisons, Figure 1B). Average temperature time-series across functional activation types exhibited slow onset dynamics (Figure 1C, black traces) that appeared to more closely resemble those of Hbt (Figure 1C, light blue traces). In contrast, CBF displayed faster onset dynamics (Figure 1C, red traces) that was also reflected in early changes in CMRO_2_ (Figure 1C, dark blue trace), since such estimates rely explicitly on CBF (Equation 1 in Methods). Notably, large temperature increases were observed during graded hypercapnia, despite the fact that CMRO_2_ decreased during evolution of the gas-challenge (Figure 1C, middle panels, black versus dark blue traces). Taken together, these results suggest that temperature increases during different forms of brain activation in our rodent model are primarily hemodynamic dependent.

**Figure 1:**
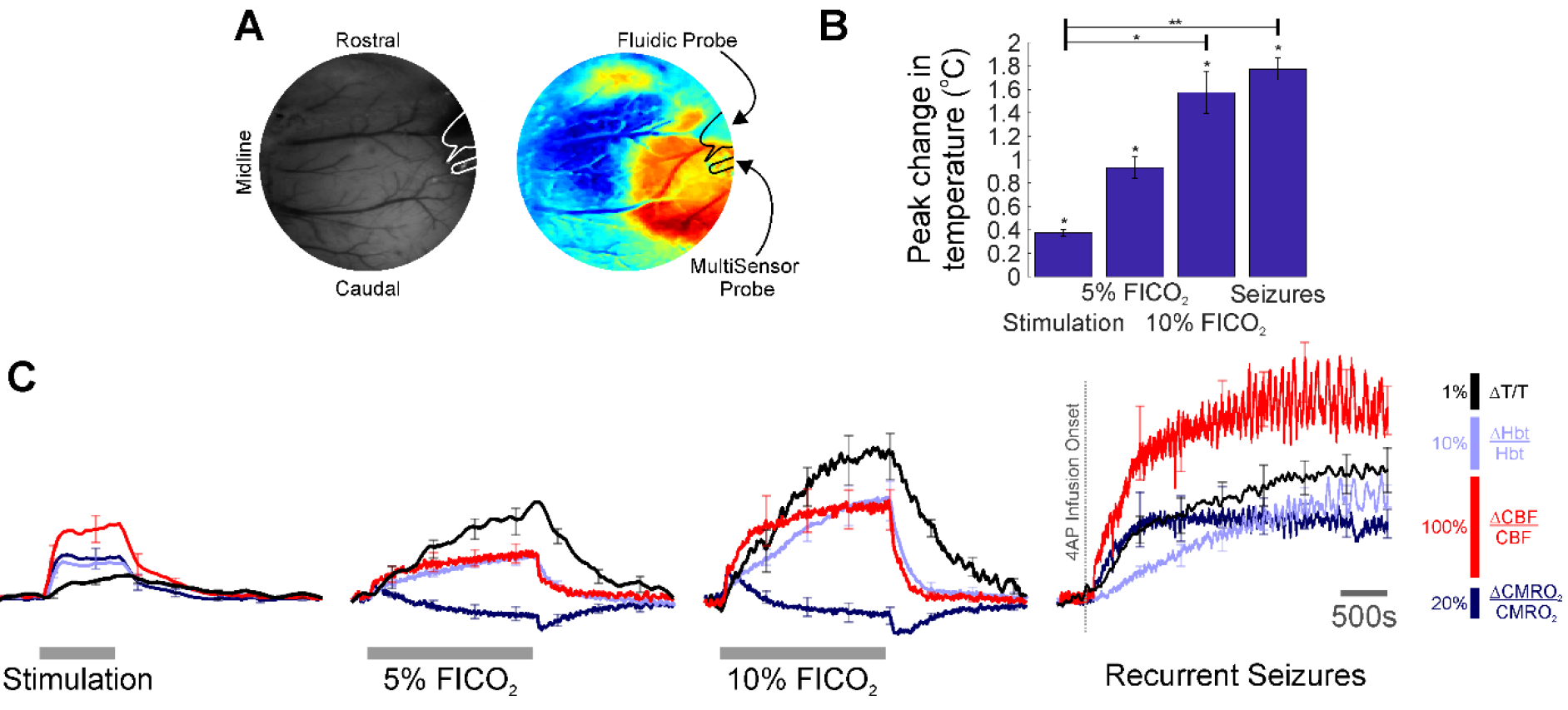
Temperature, hemodynamic and oxidative metabolism changes during sensory stimulation, graded hypercapnia and recurrent acute neocortical seizures. A) Digital image of cortical surface showing implantation sites of recording probes in a representative animal (left), colocalised to barrel cortex activated by whisker stimulation, and visualized using SPM (right, blue and red colors indicate negative and positive activation areas, respectively). B) Averaged peak change in temperature during whisker stimulation (N=5), 5% and 10% fraction of inspired CO_2_ (FICO_2_, N=5 in each case) and recurrent seizures (N=6). Individual comparisons made using 1-tailed Wilcoxon signed-ranks tests. Statistical comparisons made using Kruskal-Wallis tests with Tukey-Kramer correction. *p<0.05 and **p<0.01. C) Percentage change from baseline in cortical temperature (black), Hbt (light blue), CBF (red) and CMRO_2_ (dark blue) during sensory stimulation (left panel), graded hypercapnia (middle panels) and recurrent acute neocortical seizures (right panel, first 2500s of seizure activity), averaged across animals. Error bars are SEM.

To investigate this further, we conducted trial-by-trial analysis to assess the role of oxidative metabolism and perfusion-related variables on the observed temperature changes across experimental conditions. Weak and variable correlations were observed during sensory stimulation, most probably due the small induced changes in cortical temperature leading to a reduced signal to noise ratio (Figure 2, top panels). However, following automated removal of three temperature outliers (defined as > 3 scaled median absolute deviations away from the median, Figure 2, bottom panels, red crosses), the most prominent finding was that of a weak correlation between cortical temperature and Hbt with borderline significance (r=0.3, p=0.05, false-discovery rate (FDR) corrected for multiple comparisons, N=72, Figure 2, top middle panel). In contrast, during hypercapnia, robust and significant linear correlations were observed between temperature and Hbt (r=0.75, p=2.3e^−6^, FDR corrected for multiple comparisons, N=30), and temperature and CBF (r=0.72, p=7.6e^−6^, FDR corrected for multiple comparisons, N=30) (Figure 2, middle panels). Finally, with respect to recurrent seizures, strong linear correlations between cortical temperature changes and CMRO_2_ (r=0.65), Hbt (r=0.99), and CBF (r=0.54), were observed (Figure 2, bottom panels), but only reached significance in the case of Hbt (p=5.3e^−5^, FDR corrected for multiple comparisons, N=6).

**Figure 2:**
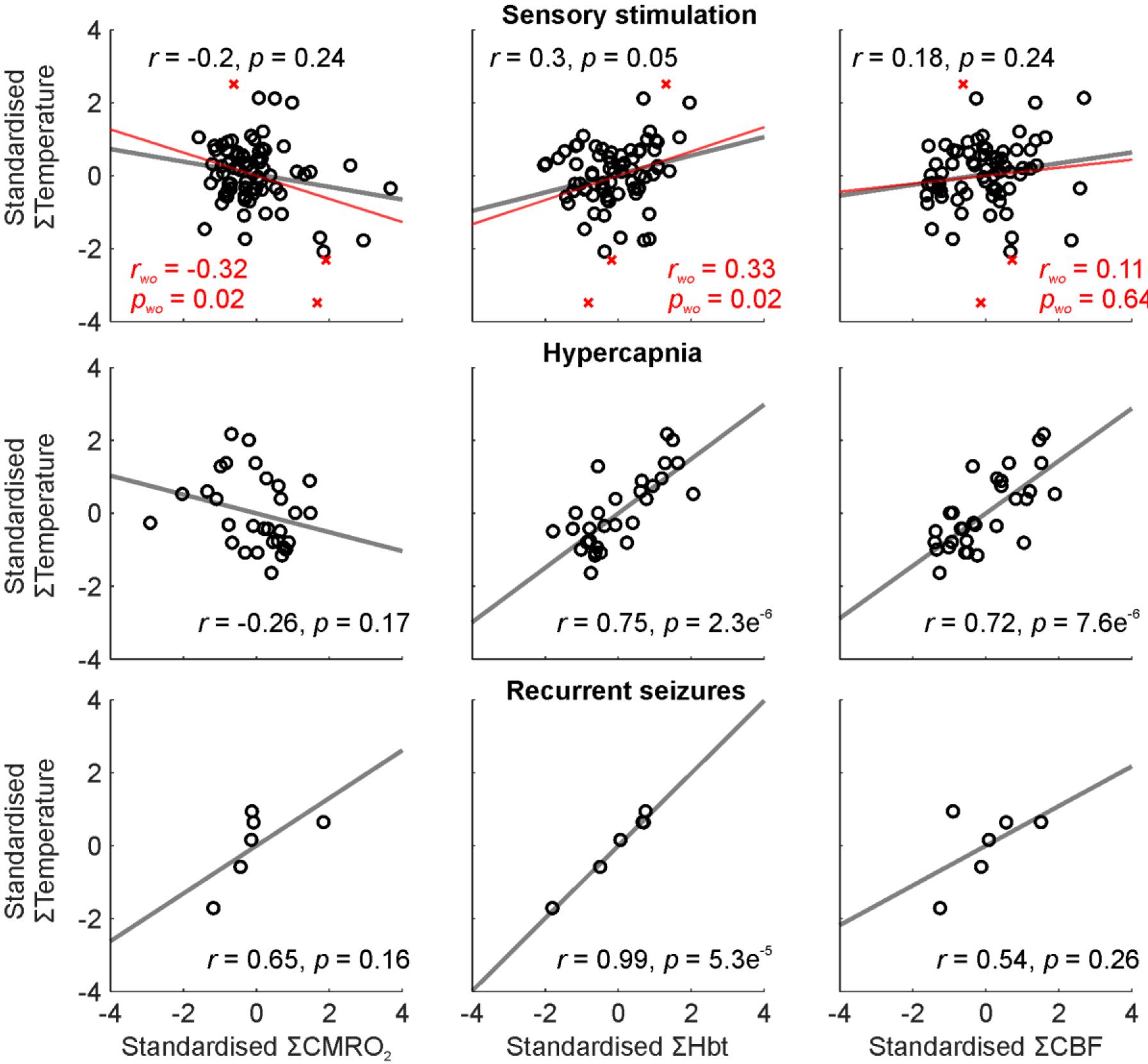
Relationship between temperature and CMRO_2_, Hbt and CBF during a range of brain activation types. Trial-by-trial analysis of coupling between standardized CMRO_2_ (left), Hbt (middle) and CBF (right), and temperature, during sensory stimulation (top, N=72), pooled hypercapnia (middle, N=30), and recurrent seizures (top, N=6). Linear functions (grey) fitted using least squares regression with Pearson correlation coefficients and significance given as insets in each case. Red regression lines in the case of sensory stimulation (top) denote fit inclusive of disregarded outliers (N=3, red crosses) with associated correlation coefficient and p-value (red text).

In combination, these results indicate that the observed cortical temperature changes in our anesthetized rodent model are predominantly perfusion-dependent, and particularly reflect changes in Hbt, across a range of brain activation types.

### Temperature increases during brain activation can modulate BOLD fMRI signal intensity

We next assessed whether the observed temperature changes recorded during brain activation could, in of themselves, affect BOLD fMRI signal intensity, using a saline-based phantom and a fast low-angle shot (FLASH) fMRI approach (Figure 3A). Relaxation of the pre-heated phantom from 38-24°C produced a ~38% increase in BOLD signal intensity across all TEs and TRs examined (Figure 3C-D, left panels). Importantly, this increase in signal intensity was not due to time/repeated scanning, since signal intensity remained approximately constant during recurrent acquisitions over a similar period of time at room temperature (Figure 3B).

The relationship between phantom temperature and signal intensity in each scanning condition was well described using a decreasing second-order polynomial function (R^2^=0.99 in all three cases, light blue text, Figure 3C-D right panels). This inverse coupling was preserved across all examined TE and TR combinations (TE/TR=6/125ms, 12/125ms, 6/1000ms), such that a generalized second-order polynomial with averaged coefficients from each condition’s individual model provided an excellent fit to the data (R^2^=0.96-0.99, light red text, Figure 3C-D right panels). This indicates that the inverse relationship seen between temperature and BOLD signal intensity appears to be independent of T1 or T2* weighting, and that the observed change in signal intensity arises due to the intrinsic influence of temperature on proton spin-density effects.

**Figure 3:**
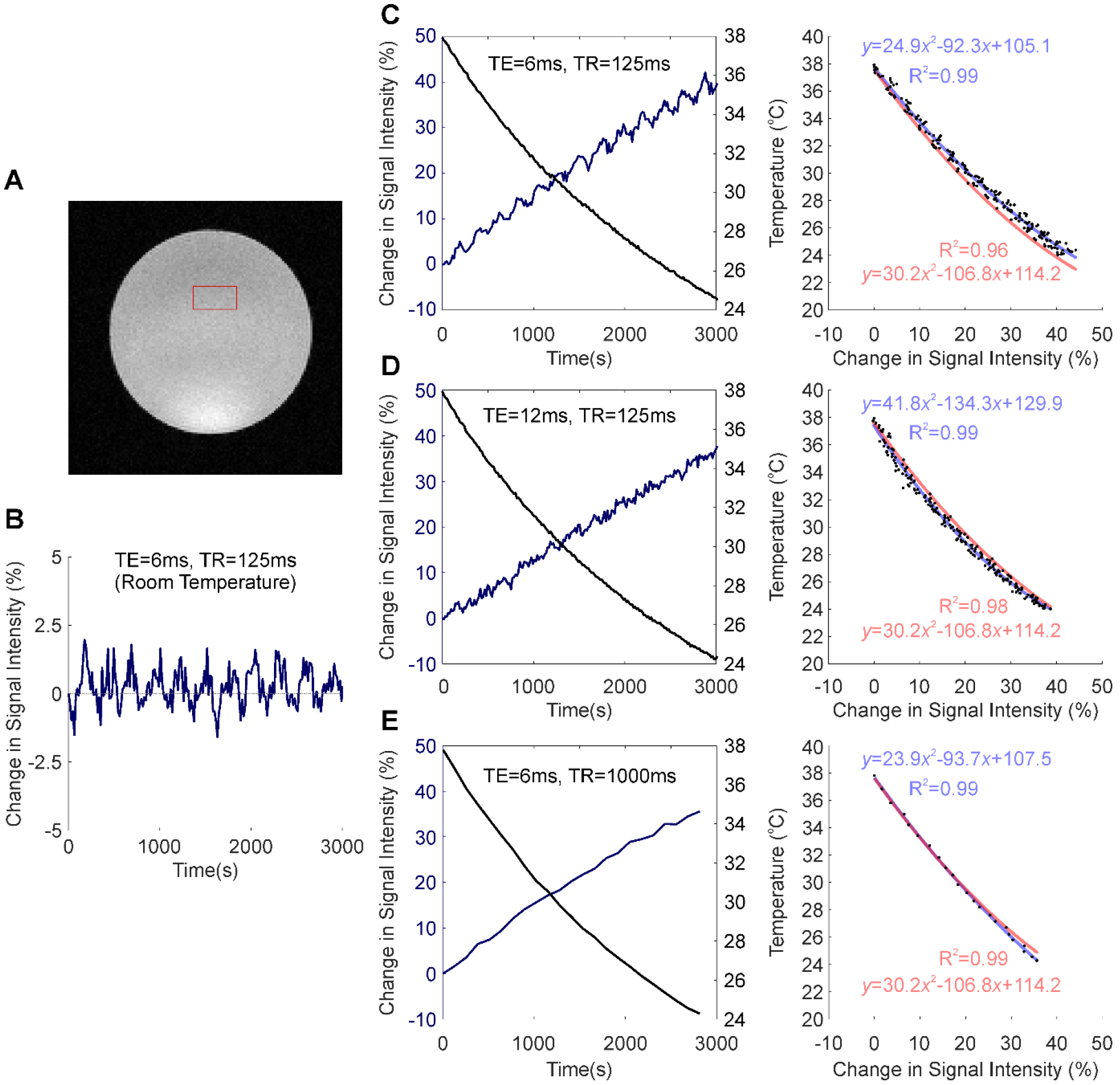
Temperature-dependence of the BOLD fMRI signal. A) Coronal image of saline-phantom, with inset of region of interest (ROI) from which fMRI timeseries data was extracted. B) Confirmation that BOLD signal intensity remains approximately constant at room temperature during repeated scan acquisitions over 3000s (TE/TR=6/125ms). C) Left: Percentage change in BOLD signal intensity during phantom-cooling from 38-24°C, TE/TR=6/125ms. Right: Associated inverse coupling between BOLD signal intensity and temperature from data shown in left panel. Second order polynomial function fitted using least-squares regression (blue), which provided an excellent fit to the data (R^2^=0.99). D and E) Same as in C, but with TE/TR=12/125ms and 6/1000ms, respectively. C-E) Generalized second-order polynomial functions fitted using least-squares and averaged coefficients from individual regression models in C-E (red) provided comparable fits (R^2^=0.96-0.99).

We subsequently employed the generalized second-order model to estimate what effect the observed activation-dependent temperature increases might have on BOLD signal intensity, during in-vivo neuroimaging. Based on this model, a 0.4°C increase in cortical temperature due to functional hyperemia, from a baseline of 32.4°C, would be associated with a ~1% decrease in BOLD signal intensity (Figure 4, light blue). 5% and 10% hypercapnia, in turn, with increases of 0.9 and 1.6°C from baseline, would equate to an approximate reduction in BOLD signal intensity of 2.2% and 4%, respectively (Figure 4, light and dark green). Finally, the dramatic 1.8°C increase in temperature seen during recurrent seizures would likely be accompanied by a 4.4% decrease in BOLD signal (Figure 4, blue). Taken together, these estimates suggest that hemodynamic-dependent increases in cortical temperature have the potential to markedly confound BOLD neuroimaging data and the modelling of such signals in anesthetized rodents.

**Figure 4:**
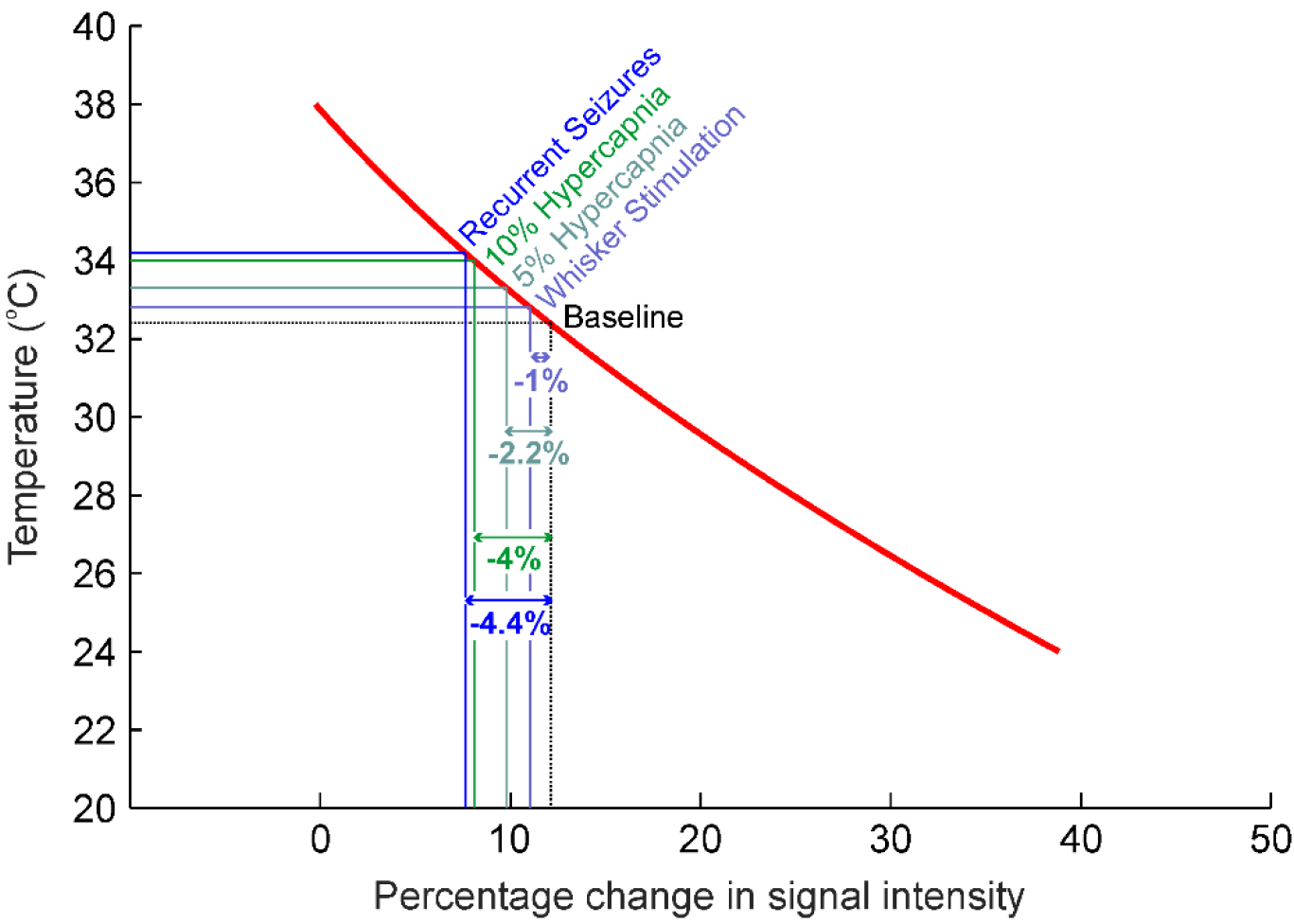
Estimates of percentage changes in BOLD signal intensity due to intrinsic temperature variations such as those observed during sensory stimulation, graded hypercapnia and recurrent seizure activity (Figure 1B). Red regression curve from generalized regression model obtained in Figure 4.

## Discussion

The current study focused on using a novel methodology to characterize temperature changes, and their underpinning hemodynamic-metabolic drivers, during a range of physiological and pathological brain activation in a popular anesthetized rat preparation. Of note, we show there to be significant increases in cortical temperature during stimulation, hypercapnia, and recurrent seizures, that were preferentially correlated to changes in total hemoglobin concentration (i.e. cerebral blood volume). Furthermore, using phantom-based evaluation, we reveal that such temperature increases *per se* can induce marked decreases in BOLD fMRI signal intensity. These findings confirm and extend previous investigations into the etiology and dynamics of cortical temperature changes during functional activation in anesthetized rodents, and have important implications to the interpretation and calibration of perfusion-related neuroimaging signals in pre-clinical models.

### Significant increases in cortical temperature during a spectrum of brain activation are Hbt-dependent

We detected significant increases in cortical temperature during whisker stimulation (0.4 °C), hypercapnia (5%, 0.9°C; 10%, 1.6°C) and recurrent seizures (1.8 °C). Previous work examining brain temperature changes in small animal models have yielded variable results, both within and across different forms of brain activation. Thermocouple based invasive measures of temperature changes to differing stimuli have been reported as 0.001-0.25°C in the anesthetized and awake cat (McElligott and Melzack, 1967; Melzack and Casey, 1967), and ~0.1°C in the chloral hydrate/halothane anesthetized rat (Lamanna et al., 1989). Other studies using a multi-sensor probe, as used here, also reported a ~0.1°C increase during long-duration forepaw stimulation in the alpha-chloralose anesthetized rat (Trübel et al., 2006). More recently, infrared imaging of rat somatosensory cortex revealed significant increases ranging from 0.05-0.1°C during whisker stimulation in the urethane-anesthetized rat (Suzuki et al., 2012). Similar variability has also been seen under hypercapnia, where global increases in CBF are induced, in which increases in brain temperature in the range 0.1-0.9°C have been previously reported in rat under different anesthetic regimes (Lamanna et al., 1989; Trübel et al., 2004; Zhu et al., 2009). So too also in anesthetized rat models of experimental epilepsy, such as >0.25°C (Lamanna et al., 1989), 0.3°C (Yang et al., 2002), and 1.2°C (Trübel et al., 2004), during induced pentylenetetrazol, 4-AP and bicuculline seizures, respectively.

The cause of this variability in the literature is most likely driven by differences in the brain/body-core temperature differential across methodologies, which are invariably negative in small animals that possess brains with large surface-to-volume ratios that are susceptible to increased heat loss to the environment. Functional hyperemia in such models, where brain temperature is cooler than the core, thus induces cerebral temperature to increase and approach that of the incoming core-temperature blood through heat-exchange (Collins et al., 2004
; Sukstanskii and Yablonskiy, 2006; Zhu et al., 2009). As a result, differences in brain exposure to the environment during experiments (Collins et al., 2004; Kalmbach and Waters, 2012) as well as anesthetic regime, through anesthesia-dependent influences of resting blood flow (Zhu et al., 2004; Zhu et al., 2009), can substantially affect the degree of negative brain/body-core temperature differential and underpin inconsistency in the aforementioned reports. In contrast, in humans and other large animals, brain temperature may be elevated relative to the core (i.e. positive brain/core temperature differential), such that functional hyperemia acts as a coolant and can reduce cerebral temperature by ~0.2°C during prolonged visual stimulation in human (Yablonskiy et al., 2000). Heat production as a byproduct of increased metabolic rate also contributes to cerebral temperature changes during functional activation, but is thought to play only a marginal role compared to perfusion changes in the presence of large brain-core temperature differentials (Yablonskiy et al., 2000; Collins et al., 2004; Zhu et al., 2006; Zhu et al., 2009).

Our results therefore confirm and extend the above studies in that temperature changes across experimental conditions were positively coupled to hemodynamic changes, and importantly, significantly correlated to those of Hbt, with little or no dependence on changes in CMRO_2_ (Figure 2). This tendency can be explained by the fact that temperature changes were slow to peak, in a similar manner to that of Hbt and in contrast to CBF. Furthermore, given that capillary hyperemia may underpin delayed Hbt dynamics (Hillman et al., 2007), and the propensity for heat transfer at the capillary level through extensive fluid exchange (Yablonskiy et al., 2000), we suggest that Hbt changes in the capillary bed underpins the observed temperature changes. This would also follow under the presumption that Hbt is biased to changes in capillary regions, due to the ROI capturing primarily the parenchyma with relatively less surface microvasculature. Notably, this finding indicates that measures of Hbt/CBV could be valuable inputs to theoretical models of brain temperature (Yablonskiy et al., 2000; Collins et al., 2004; Sotero and Iturria-Medina, 2011).

### Temperature-dependent effects on the BOLD fMRI signal

Cerebral temperature changes are known to affect metabolic rate (Swan, 1974) and the affinity of hemoglobin for oxygen (Hall, 2015). Temperature fluctuations thus influence the blood BOLD fMRI signal, which relies on the poorly understood coupling between cerebral blood flow, metabolism and hemoglobin oxygenation (Ogawa et al., 1993). Indeed, recent theoretical work has established the relationship between changes in temperature and the BOLD fMRI signal in the human brain (Yablonskiy et al., 2000; Sotero and Iturria-Medina, 2011). Manipulated increases in anesthetized rat body temperature have also been demonstrated to decrease the BOLD fMRI signal, a phenomenon attributed to decreased affinity of hemoglobin for oxygen and an increase in deoxyhemoglobin concentration (Vanhoutte et al., 2006). However, MR relaxation properties have also been shown to relate to manipulated temperature of ex-vivo human tissue samples (Petrén-Mallmin et al., 1993). This implies that the BOLD fMRI signal can be modulated by temperature changes, in of themselves, even in the absence of physiological activation and the subsequent temperature-dependent modulation of hemodynamic-metabolic processes that underpin the BOLD signal in-vivo. Surprisingly, this possibility has not been previously investigated, and is of particular relevance to anesthetized rodent BOLD neuroimaging studies where functional activation induces significant increases in cortical temperature, as described above. Here, using a phantom-based evaluation, we demonstrate that temperature increases, such as those observed during whisker stimulation, hypercapnia and recurrent seizures in our model (0.4-1.8°C), markedly decrease BOLD signal intensity by ~1-4% (Figure 4) with a fast FLASH based MR sequence. We elected not to use a standard Echo-Planar Imaging (EPI) approach to assess the BOLD signal change as a function of temperature since an EPI pulse sequence has a confounding signal drift caused by the high imaging readout load on the system (Vos et al., 2017). This causes heating effects, both in the sample and electronics, which can decrease, for example, fat suppression efficiency (Thesen et al., 2003), and decrease the scanner’s temporal stability (Benner et al., 2006). Disambiguating the sample temperature dependent signal from the above would therefore have proved inherently more challenging. We were able to bypass these potential signal confounds using a FLASH based Gradient Echo sequence at a lower image temporal resolution, but same image/relaxation contrast, since the timeframe for the phantom to decrease in temperature was approximately 3000s. This intrinsic inverse effect of temperature on BOLD signal intensity, as well as the increased metabolic rates and facilitated unloading of oxygen from hemoglobin as a result of activation-induced temperature increases, such as those presented here, should therefore be taken into consideration when interpreting BOLD fMRI signal changes in small animal models that possess negative brain-core temperature differentials, and which can be exacerbated by degree of brain exposure and anesthesia. This is particularly relevant to the growing number of studies examining the neurophysiological underpinnings of the BOLD signal using simultaneous BOLD fMRI and optical modalities, such as calcium imaging (Schulz et al., 2012; Liang et al., 2017) and optogenetics (Lee et al., 2010; Yu et al., 2016; Schmid et al., 2017), that are typically conducted using anaesthetized and anesthetized rodent models (including the model described here), and require a craniotomy for imaging and/or illumination. It is also pertinent to studies using hypercapnia to calibrate fMRI signals to a baseline CMRO_2_ and examine subsequent transients during functional activation (Davis et al., 1998; Kida et al., 2007).

## Conclusion

In conclusion, this study provides novel insights into the dynamics and hemodynamic drivers of cortical temperature changes during a range of physiological and pathological brain activations in a commonly used anesthetized rat model, and urges caution when calibrating and interpreting BOLD signals in preclinical models where temperature increases during gas challenges and functional activation are likely to have a confounding effect. Further elucidating the feedback relationship between cortical temperature and hemodynamic and metabolic processes during functional activation, in the context of brain-core temperature differentials, and subsequent (and intrinsic temperature) effects on the BOLD signal, will require tight control of brain temperature alongside invasive measures of hemodynamic and oxidative metabolism, and concurrent BOLD fMRI imaging. This is currently in development in our laboratory, and will be a crucial next step to understanding the role of brain temperature variations in regulating cerebral processes, and the impact of these on existing and novel functional neuroimaging signals in preclinical models.

## Acknowledgements

We thank the technical staff of the University of Sheffield’s Department of Psychology, Natalie Kennerley and Michael Port.

